# Predicting transcription factor activity using prior biological information

**DOI:** 10.1101/2022.12.16.520295

**Authors:** Joseph Estabrook, William M. Yashar, Hannah D. Holly, Julia Somers, Olga Nikolova, Özgün Barbur, Theodore P. Braun, Emek Demir

## Abstract

Transcription factors are critical regulators of cellular gene expression programs. Disruption of normal transcription factor regulation is associated with a broad range of diseases. In order to understand the mechanisms that underly disease pathogenesis, it is critical to detect aberrant transcription factor activity. We have developed Priori, a computational method to predict transcription factor activity from RNA sequencing data. Priori has several key advantages over existing methods. Priori utilizes literature-supported regulatory relationship information to identify known transcription factor target genes. Using these transcriptional relationships, Priori uses linear models to determine the impact and direction of transcription factor regulation on the expression of its target genes. In our work, we evaluated the ability of Priori and 16 other methods to detect aberrant activity from 124 single-gene perturbation experiments. We show that Priori identifies perturbed transcription factors with greater sensitivity and specificity than other methods. Furthermore, our work demonstrates that Priori can be used to discover significant determinants of survival in breast cancer as well as identify mediators of drug response in leukemia from primary patient samples.

## INTRODUCTION

Dysregulation of transcription factor regulatory networks is a common driver of disease^1,2^. Transcription factors establish and maintain cellular gene expression programs by binding to *cis*-regulatory DNA sequences and recruiting cofactors that aid with gene transcription^3,4^. Many diseases are caused by mutations in these putative regulatory elements as well as in the transcription factors and co-factors that interact with them^2^. In cancer, over-expression of oncogenic transcription factors contributes to tumorigenesis through altered expression of downstream target genes^5–7^. Therefore, detection of abnormal transcription factor activity is valuable for better understanding the mechanisms that underly disease pathogenesis.

Gene expression profiling, including RNA sequencing (RNA-seq), is commonly used to monitor dynamic changes in transcription factors and their gene regulatory networks. Initial studies to infer transcription factor activity used gene expression as a proxy for activity^8–10^. However, this approach has several shortcomings. Gene expression is only an indirect measurement of protein activity due to the complex mechanisms controlling protein synthesis and degradation^11–13^. Feedback loops may alter the expression of transcription factors in response to their regulatory activity^2,14–16^. Moreover, transcription factors that play a central role in coordinating the active gene transcription program tend to be lowly expressed and are rarely differentially expressed^17,18^. Reliable predictions of transcription factor activity, therefore, cannot be limited to evaluating transcription factor expression alone.

An alternative approach to inferring transcription factor activity is to assess the expression of their downstream target genes^8–10^. This approach has two major benefits. First, evaluation of hundreds or thousands of downstream targets instead of a single transcription factor likely improves the prediction robustness. While some of these targets may be context-specific, analyzing them in aggregate likely improves the prediction generalizability across many contexts. Second, as target gene expression is downstream of transcription factor control, these signatures are expected to reflect the actual transcriptional impact more accurately. Therefore, accounting for the downstream impact of transcription factors on its gene regulatory networks is important for activity inference.

Multiple methods have been developed to quantify transcription factor activity from gene expression data. These approaches can be grouped based on how they select gene expression features. Methods like Univariate Linear Model (ULM) and Multivariate Linear Model (MLM) use every gene in a dataset, nominating transcription factors as a covariate that best estimates the expression of all other genes^19^. However, these methods develop activity signatures using genes that may not have a true biological relationship to the transcription factor of interest. Gene set approaches like Over Representation Analysis (ORA), Fast Gene Set Enrichment (FGSEA), Gene Set Variation Analysis (GVSA), and AUCell infer activity using sets of published transcription factor target genes or target genes curated by experts^20–23^. While gene set methodologies are simple and popular, they are susceptible to the quality and comparability of gene set signatures^22^. Network inference approaches, including Algorithm for the Reconstruction of Accurate Cellular Networks (ARACNe), infer gene regulatory networks based on the covariance of transcription factors and its putative targets^24,25^. The same group that developed ARACNe also created Virtual Inference of Protein-activity by Enriched Regulon analysis (VIPER) to infer transcription factor activity from ARACNe gene expression signatures^26^. The challenge with these approaches is deconvoluting pleiotropy, where the expression of a target gene is controlled by multiple transcription factors^2^. While some of these methods, including VIPER, has an option to correct for pleiotropy, the problem of network inference often remains underdetermined as there are many possible solutions that can explain the underlying data^8,27^. Although other methodologies have constrained these networks using biological data generated from orthogonal modalities, grounding these predictions to actual transcription factor activity remains challenging^27–36^.

Here, we propose an approach that uses prior, peer-reviewed biological information to infer transcription factor activity called Priori. Our method has two major advantages over the existing methods. First, Priori identifies transcription factor target genes using carefully extracted transcriptional regulatory networks from Pathway Commons^37,38^. This resource continually collects information on biological pathways including molecular interactions, signaling pathways, regulatory networks, and DNA binding. Pathway Commons currently contains data from 22 high-quality databases with over 5,700 detailed pathways and 2.4 million interactions. Second, using the transcriptional relationships from Pathway Commons, Priori fits linear models to the expression of transcription factors and their target genes. These models allow Priori to understand the impact and direction of transcription factor regulation on its known target genes. Our findings show that Priori detects aberrant transcription factor activity with greater sensitivity and specificity than other methods. Furthermore, we demonstrate that Priori can be deployed to discover significant predictors of survival in breast cancer as well as identify mediators of drug response in leukemia from primary patient samples.

## RESULTS

### Priori uses prior biological information to infer transcription factor activity

For each transcription factor in an RNA-seq dataset, Priori generates an activity score (Figure 1a). Priori first identifies the known target genes for each transcription factor in a dataset from Pathway Commons (or another network provided by the user)^37,38^. Priori then assigns weights to each target gene by correlating the target gene expression to its transcription factor. To identify the targets that are most impacted by transcription factor regulation, Priori separates the up- and down-regulated genes by their transcription factor-target gene weights and ranks their expression. These ranks are subsequently scaled by the transcription factor-target gene weights. The activity score for each transcription factor is then calculated by summing the weighted ranks. With this single-component model, researchers can use Priori to predict transcription factor activity from RNA-seq data.

**Figure 1:**
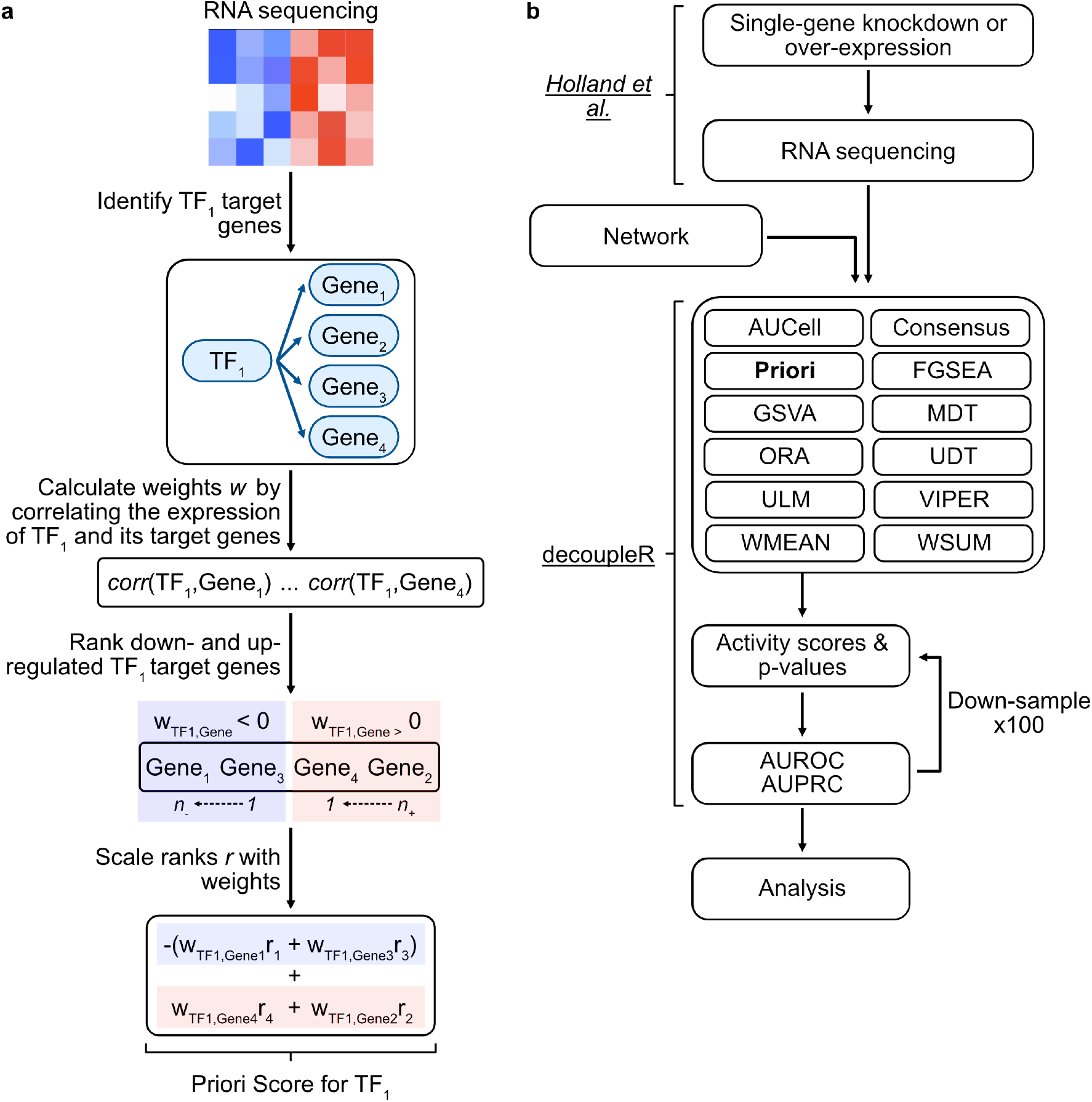
Overview of the Priori methodology and benchmarking workflow. a. Priori generates an activity score for each transcription factor in an RNA-seq dataset. Priori first extracts the downstream target genes for each transcription factor from Pathway Commons. Priori then calculates weights for each target gene by correlating the expression of each transcription factor to its target genes. Priori then ranks the absolute expression of all genes in the dataset and scales these ranks by the transcription factor-target gene weights. The summation of the weighted target gene ranks is the transcription factor activity score. b. Schematic overview of the benchmarking workflow. We generated transcription factor activity scores for each method using normalized RNA-seq counts following single-gene knockdown or over-expression. Priori, along with other methods that use prior information, generated activity scores with transcriptional relationships from Pathway Commons. AUROC and AUPRC values were calculated for each down-sampling permutation. 100 down-sampling permutations were performed to compare an equal number of perturbed and unperturbed genes.

In order to compare how well Priori detects aberrant transcription factor activity to other methods, we used the decoupleR benchmarking workflow (Figure 1b)^39^. The decoupleR workflow facilitated a common evaluation scheme to determine how often each method correctly identified transcription factors that have been knocked-down or over-expressed from RNA-seq data^40^. Using this workflow, we generated transcription factor activity scores for 17 methods, including Priori, using the default parameters. Several of these methods were developed by the authors of the decoupleR workflow, including the standard, normalized (Norm), and corrected (Corr) versions of Weighted Mean (WMEAN), Weighted Sum (WSUM), Univariate Decision Tree (UDT), and Multivariate Decision Trees (MDT)^39^. For methods that use prior information, we generated activity scores using the Pathway Commons transcriptional relationships. We ranked the transcription factor activity scores for each experiment and compared the ranks of perturbed and unperturbed transcription factors. We evaluated how often the perturbed transcription factor activity score was among the top activity scores in each experiment. Since the number of unperturbed transcription factors vastly outnumbered the perturbed transcription factors, we implemented a down-sampling strategy. For each down-sampling permutation, we calculated area under the precision recall curve (AUPRC) and receiver operating characteristic (AUROC) metrics. The results of the decoupleR workflow allowed us to compare the sensitivity and specificity of Priori to detect perturbed transcription factor activity to 16 other methods.

### Priori identifies perturbed transcription factors with greater sensitivity and specificity than other methods

We generated activity scores for 124 knockdown and over-expression experiments across 62 transcription factors (94 knockdown and 30 over-expression experiments; Supplementary Table 1). To understand the patterns in predicted activity across the methods, we correlated the activity scores (Figure 2a). The Priori activity scores were dissimilar from the other methods, indicating that Priori identified a unique pattern of transcription factor activity. To assess how well the methods detected perturbed transcription factors, we calculated AUPRC and AUROC metrics for each down-sampling permutation (Figure 2b-d). Priori had a greater AUPRC and AUROC values than the other methods across all experiments (Supplementary Table 2). ULM, Norm WMEAN, Norm WSUM, and VIPER were the next closest methods by AUPRC and AUROC. Collectively, these experiments show that Priori detects perturbed transcription factors with greater sensitivity and specificity than other methods.

**Figure 2:**
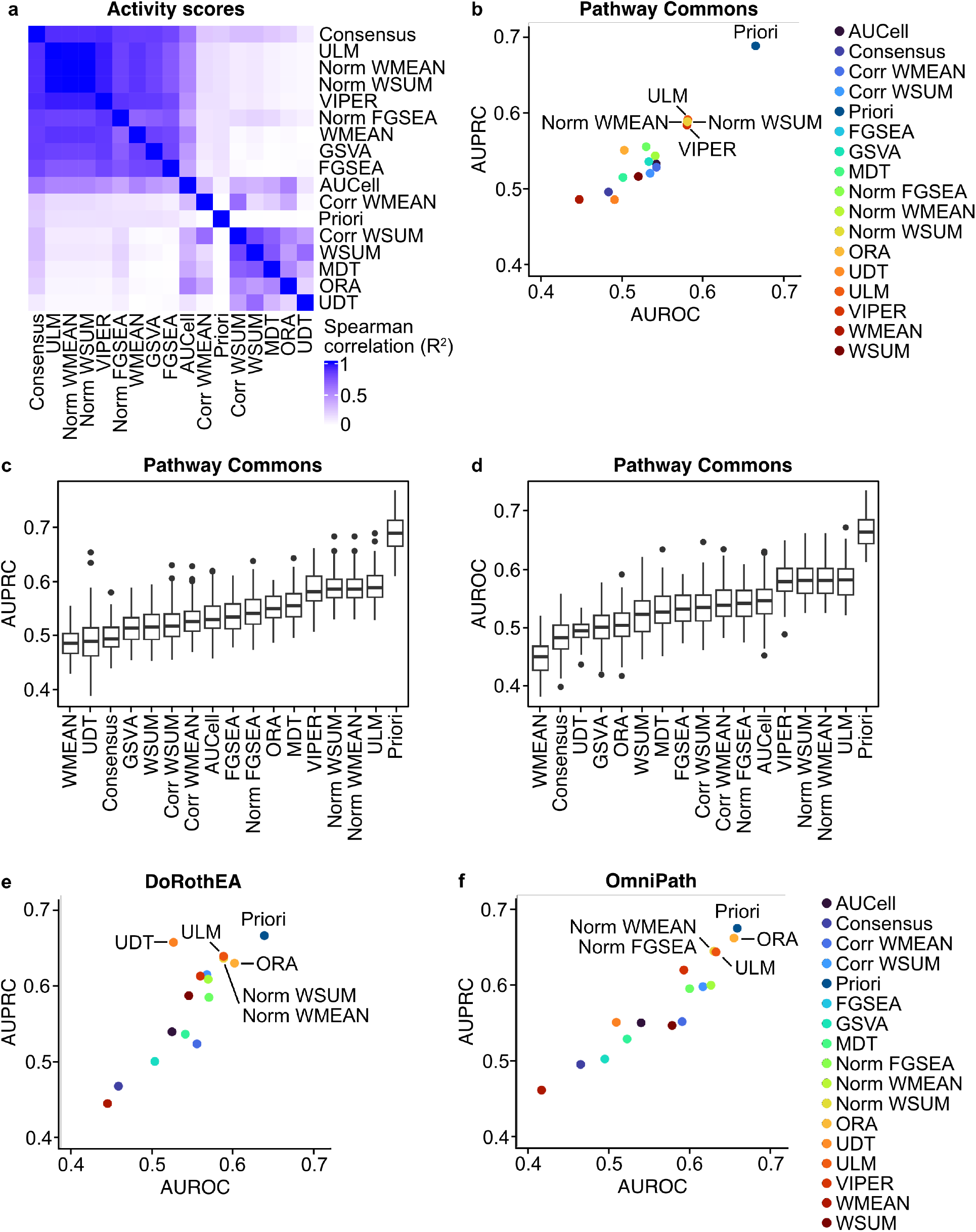
Priori detects aberrant transcription factor activity with improved sensitivity and specificity. a. Using the decoupleR workflow, transcription factor activity scores were generated using the perturbation dataset. Spearman correlation of the activity scores for each method. b. Activity scores were generated using Pathway Commons transcriptional relationships. Mean AUPRC and AUROC values across the 100 down-sampling permutations for each method. c, d. The distribution of (c) AUPRC and (d) AUROC values across the 100 down-sampling permutations from (b). e, f. Activity scores were generated using (e) DoRothEA or (f) OmniPath transcriptional relationships. Mean AUPRC and AUROC values across the 100 down-sampling permutations for each method.

While our analyses showed that Priori can identify perturbed transcription factors from RNA-seq data, we wanted to understand the extent to which prior information impacted its performance. Pathway Commons is one of several large databases that curates transcriptional relationships. DoRothEA is a comprehensive resource, which assembles transcription factor-target gene relationships from ChIP-seq peaks, inferred regulatory networks (including ARACNe), transcription factor binding motifs, and literature-curated resources^41^. OmniPath, on the other hand, integrates intra- and inter-cellular signaling networks in addition to transcriptional relationships from 100 different resources, including DoRothEA and Pathway Commons^42^. To evaluate the impact of prior information on each method, we used the decoupleR workflow to generate activity. However, we used the transcriptional relationships from DoRothEA or OmniPath instead of Pathway Commons as prior information. We found that in both instances, Priori had higher AUPRC and AUROC values than the other methods (Figure 2e, f; Supplementary Figure 1; Supplementary Table 3, 4). Priori exhibited similar AUPRC and AUROC values when using transcriptional relationships from DoRothEA, OmniPath, or Pathway Commons. However, Priori had the highest mean AUPRC and AUROC values using the Pathway Commons transcriptional relationships. Our analyses demonstrated that regardless of the prior transcriptional relationship information, Priori detected perturbed transcription factor activity with improved sensitivity and specificity.

We evaluated how transcriptional relationships from Pathway Commons, DoRothEA, and OmniPath affected the performance of each method. However, these databases do not provide the appropriate information for some of the methods, including VIPER, that were designed to generate activity scores from networks with signed edges that are weighted by likelihood. While decoupleR does not allow for networks with signed edges, we wanted to understand how VIPER’s performance changed with ARACNe-generated networks with likelihood-weighted edges^39^. The authors of ARACNe have created and released a variety of cancer-type-specific networks trained on TCGA RNA-seq datasets^25^. We identified the cell lines as well as the cancer types and organ sites associated with each cell line tested in the perturbation dataset (Supplementary Figure 2a-c; Supplementary Table 5). With decoupleR, we generated VIPER activity scores using any TCGA ARACNe network that was trained on a cancer type evaluated in the perturbation dataset. We observed that VIPER had greater AUPRC and AUROC values when using the transcriptional relationships from Pathway Commons than those from the TCGA ARACNe networks (Supplementary Figure 2d). ARACNe can also be used to reverse engineer gene regulatory networks from a RNA-seq dataset^25^. Using the new implementation of ARACNe, ARACNe-AP, we inferred transcriptional relationships using the perturbation RNA-seq data. We observed improved AUPRC and AUROC values when VIPER used ARACNe-AP transcriptional relationships as prior information (Supplementary Figure 2e). However, these AUPRC and AUROC values were still not greater than Priori using prior transcriptional relationships from Pathway Commons.

### Priori’s improved performance is due to evaluating the direction of transcriptional regulation

We demonstrated that Priori detects perturbed transcription factors with improved sensitivity and specificity, in particular when it used Pathway Commons transcriptional relationships as priori information. However, it was unclear what enabled Priori to detect aberrant activity better than other methods. While Pathway Commons does not provide signed relationships, we designed Priori to infer the impact and direction of transcriptional regulation on its target genes. Priori correlates the expression of transcription factors and their known target genes. Priori subsequently assigns the impact and direction of transcriptional regulation as the coefficient and sign of the spearman correlation, respectively. Using these correlations, we observed that Priori detected 2,507 more significant up-regulated transcriptional relationships than down-regulated relationships in the perturbation dataset (Figure 3a). Priori predicted that transcription factors had a significantly greater impact on its up-regulated targets than its down-regulated target genes (Figure 3b). These analyses show that there are important directions of transcription factor regulation on their target genes.

**Figure 3:**
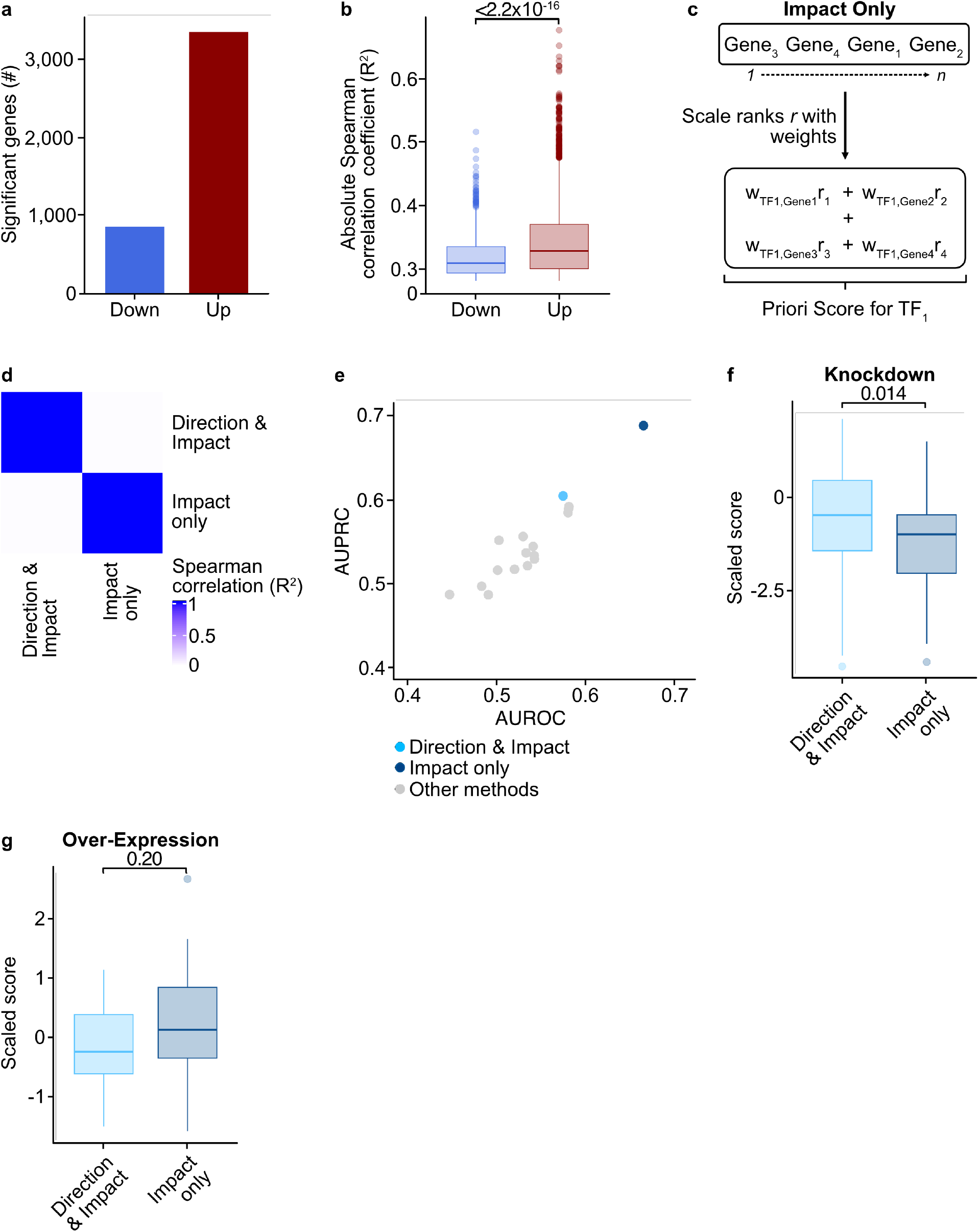
Evaluation of the direction and impact of transcriptional regulation is critical for Priori to detect aberrant transcription factor activity. a. Transcription factor target genes were identified using Pathway Commons. The expression of transcription factors and their target genes were evaluated using Spearman correlation. Statistical significance was determined using the Spearman correlation p-value with an FDR post-test correction. The Spearman correlation coefficient was used to determine down-regulated (R^2^ < 0) and up-regulated target genes (R^2^ > 0). b. Absolute Spearman correlation coefficient of the expression of transcription factors and their down-regulated or up-regulated target genes. Statistical significance was determined by a two-sided Student’s t-test. c. Schematic showing how Priori was altered to assess only the absolute impact of transcriptional regulation (in contrast to both direction and impact of regulation as depicted in Figure 1a). d. Using the decoupleR workflow, transcription factor activity scores were generated using the perturbation dataset. Spearman correlation of the activity scores for each method. e. Activity scores were generated using Pathway Commons transcriptional relationships. Mean AUPRC and AUROC values across the 100 down-sampling permutations for Priori using only the impact of transcriptional relationships. Mean AUPRC and AUROC values from the methods in Figure 2b are also shown. f, g. Priori activity scores using absolute and relative transcriptional relationships were z-transformed. Distribution of scaled scores for perturbed transcription factors across all (f) knockdown and (g) over-expression experiments. Statistical significance was determined by a two-sided Student’s t-test.

To understand how assessment of the direction of transcriptional regulation is important for detecting aberrant transcription factor activity, we altered Priori to only evaluate the absolute impact of transcriptional regulation (Figure 3c). We used the decoupleR pipeline to generate activity scores using the Pathway Commons transcriptional relationships as prior information. These activity scores were highly dissimilar to the scores generated by Priori that evaluated both the direction and impact of transcriptional regulation (R^2^ = 0.013; Figure 3d). Moreover, when Priori only uses the impact of transcriptional regulation to detect perturbed transcription factors, its AUPRC and AUROC values are less than when it uses both the direction and impact (Figure 3e). Accounting for direction of transcriptional regulation allows Priori to detect knocked down and over-expressed transcription factors. We observed an expected decrease in scaled scores across all knockdown experiments when Priori evaluates both the direction and impact of transcriptional regulation (Figure 3f). Scaled scores less than zero indicate that the activity of a transcription factor is down-regulated compared to others in the dataset (and vice versa). These scales scores are significantly less than when Priori only evaluates transcriptional regulation impact. Consistently, Priori predicted an expected increase in activity to over-expressed transcription factors when it assessed both direction and impact (Figure 3g). While these scores are not statistically different to the scores that only evaluated the impact of transcriptional regulation, the mean predicted activity from the impact-only scores was less than zero. Overall, these analyses demonstrate that assessment of the direction and impact of transcriptional regulation allows Priori to detect aberrant transcription factor activity with improved sensitivity and specificity.

### FOXA1 transcription factor activity is a significant determinant of survival among patients with breast cancer

Since we demonstrated that Priori identifies perturbed regulators with greater sensitivity and specificity than other methods, we sought to determine whether Priori could be used to understand transcription factor drivers of disease. We used Priori to generate transcription factor activity scores for 637 patients with invasive breast ductal carcinoma (BIDC)^43^. We also generated activity scores using three of the top methods from the decoupleR benchmark analysis: VIPER, ORA, and Norm WMEAN. The Priori scores from these patients clustered by breast cancer subtypes (Figure 4a; Supplementary Figure 3a). We did not observe a clear separation of breast cancer subtypes when using activity scores from the other methods (Supplementary Figures 3b-d). BIDC is classified into three molecular subtypes: luminal, HER2, and basal cancers^44–48^. The most common types, luminal and basal breast cancers, are distinguished by hormone receptor expression. Luminal breast cancers express estrogen and progesterone receptors, whereas basal breast cancers do not. Unsupervised clustering of the Priori scores revealed that the predicted activity of transcription factors that regulate the expression of estrogen receptors (*ESR1*) and progesterone receptors (*NR3C1*) were decreased in the basal breast cancer cluster (Figure 4b). Moreover, the basal breast cancer cluster had decreased predicted GATA3 activity, which is critical to luminal cell specification, as well as increased predicted MYC activity, which is consistent with previous studies^44,49–51^. Interestingly, the greatest difference in predicted activity between the two clusters was FOXA1 activity, which encodes a forkhead protein associated with mammary gland development (Figure 4c, d; Supplementary Table 6). The difference in FOXA1 activity was much less pronounced between the patient clusters defined by the other methods (Supplementary Figure 3e-g). Evaluation of the greatest difference in predicted activity between the other method clusters nominated transcription factors other than FOXA1. Specifically, this analysis nominated OR10H2 by VIPER, ACTL6A by ORA, and TGFβ2 by Norm WMEAN (Supplementary Figure 3h-j). Together, these analyses show that we were able nominate known drivers of basal and luminal breast cancer by predicting transcription factor activity with Priori. These transcription factors were distinct from those nominated by other methods.

**Figure 4:**
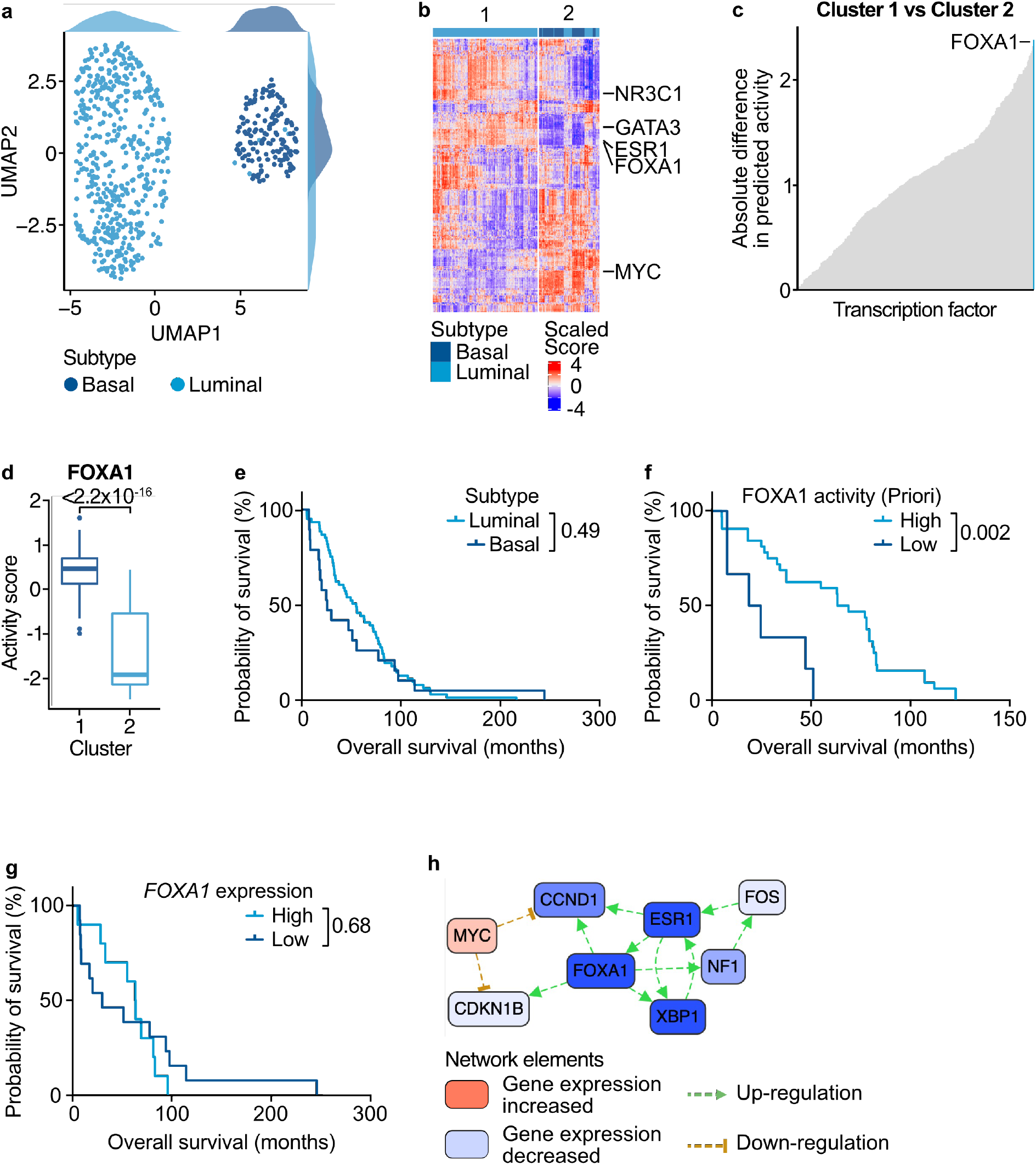
FOXA1 transcription factor activity is a significant determinant of survival for patients with BIDC. a. Priori scores were generated from RNA-seq of 637 patients with BIDC. UMAP dimensional reduction and projection of Priori scores. Dots are colored by the breast cancer molecular subtype. b. Unsupervised hierarchical clustering of Priori scores generated in (a). c. Mean absolute difference of Priori scores from patients in the clusters 1 and 2 defined in (b). d. Distribution of FOXA1 Priori scores among patients in clusters 1 and 2 defined in (b). Statistical significance was determined by a two-sided Student’s t-test. e-g. Kaplan-Meier survival analysis of patients grouped by (e) molecular subtype, (f) FOXA1 Priori scores, or (g) *FOXA1* normalized gene expression counts. Patients among the top 90% of Priori scores or counts were grouped into “High” and those in the bottom 10% were grouped into “Low”. Statistical significance was determined by a log-rank Mantel-Cox test. h. Differential gene expression network enrichment between clusters defined in (b). Select significantly enriched nodes are shown.

To understand the clinical impact of transcription factor activity in breast cancer, we evaluated survival differences among the patients in the BIDC cohort. Patients grouped by their molecular subtypes demonstrated no significant difference in survival (Figure 4e). When we grouped patients by predicted FOXA1 Priori activity scores, patients with low FOXA1 activity had a significantly decreased chance of survival (Figure 4f). Notably, we did not observe a survival difference when patients were instead grouped by *FOXA1* expression or by the FOXA1 activity scores generated by the other methods (Figure 4g; Supplementary Figure 4a-c). Additionally, we did not see a survival difference in the transcription factor drivers nominated by the other methods (Supplementary Figure 4d-f). These data suggest that FOXA1 transcription factor activity, not just expression, is a critical determinant of survival for patients with BIDC. In order to understand the critical FOXA1 target genes, we generated a differential gene regulatory network between the basal and luminal populations (Figure 4h)^37,38,52^. This network revealed that FOXA1 target genes *ESR1* and *XBP1*, which promotes basal breast cancer progression, are significantly decreased among the patients with basal breast cancer (Supplementary Table 7)^53^. These analyses nominate FOXA1 regulation of *ESR1* and *XBP1* expression as important regulators of survival in BIDC.

### FOXO1 transcription factor activity mediates venetoclax resistance in leukemia

Aberrant transcription factor activity is also an important regulator of drug resistance in multiple tumor types^1,54–57^. As we have shown that Priori nominates transcription factor regulation associated with breast cancer survival, we wanted to understand whether Priori could also be used to identify mediators of drug sensitivity. Therefore, we calculated Priori scores from RNA-seq of 859 patients with acute myeloid leukemia (AML)^58,59^. These samples were also characterized with *ex vivo* drug sensitivity data. Additionally, we generated activity scores using three of the top methods from the decoupleR benchmark analysis that were also used in the breast cancer analysis (VIPER, ORA, and Norm WMEAN). To nominate transcription factors mediators of drug sensitivity, we correlated predicted transcription factor activity scores from each method to drug response. We found 11,075 significant inhibitor-transcription factor activity relationships using Priori, 192 of which were also identified by VIPER, ORA, and Norm WMEAN (Figure 5a; Supplementary Figure 5a; Supplementary Table 8). VIPER likely identified the most significant inhibitor-transcription factor activity relationships (29,682) because it generated scores for more transcription factors than the other methods (1,363; Supplementary Figure 5b). Among the strongest correlations from Priori was predicted FOXO1 transcription factor activity with venetoclax resistance (R^2^ = −0.59; Figure 5b). Venetoclax resistance is more highly correlated with predicted FOXO1 activity than *FOXO1* expression alone (R^2^ = −0.5; Figure 5c). While FOXO1 Norm WMEAN activity scores were directly proportional to venetoclax resistance, the VIPER and ORA activity scores were anti-correlated to venetoclax resistance (Supplementary Figure 5c-e). These findings nominate FOXO1 activity as a mediator of venetoclax activity in AML.

**Figure 5:**
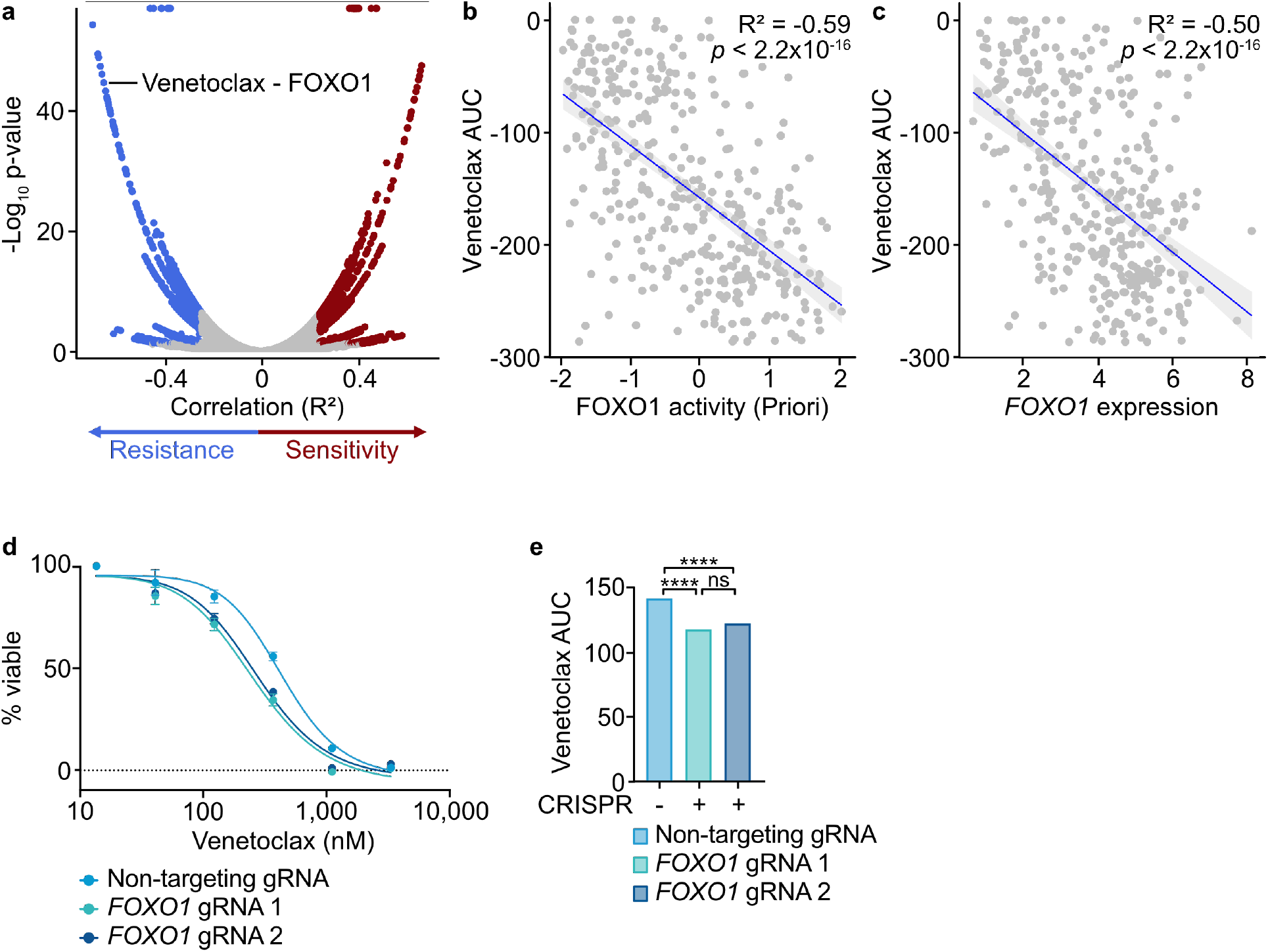
FOXO1 is a critical mediator of response to venetoclax in AML. a. Priori scores generated from RNA-seq of 859 patients with AML. Spearman correlation of Priori scores and *ex vivo* drug response AUC data. b, c. Spearman correlation of venetoclax AUC and (b) FOXO1 Priori scores or (c) *FOXO1* normalized counts. Statistical significance was determined using the Spearman correlation p-value with an FDR post-test correction. d, e. THP-1 cells were transduced with lentiviral particles harboring expression cassettes for hSpCas9 and a non-targeting or *FOXO1* guide RNA. Cells were cultured for 3 days along a 7-point curve with venetoclax. Cell viability was assessed by CellTiter Aqueous colorimetric assay. ns = not significant; * = p < 0.05, ** = p < 0.01, *** = p < 0.001, **** = p < 0.0001.

Venetoclax induces cancer cell death by restoration of intrinsic mitochondrial apoptosis. Venetoclax blocks BCL2 from sequestering factors that activate pro-apoptotic BCL2 family proteins, such as BAX^60^. In mantle cell lymphoma (MCL), it has been reported that genomic regions of BAX and multiple other pro-apoptotic BCL2 family proteins are bound by FOXO1^61^. The authors further demonstrated that disruption of FOXO1 activity sensitized MCL cell lines to venetoclax. However, the relationship between FOXO1 activity and venetoclax resistance has not been investigated in AML. Priori work has shown that monocytic AML is intrinsically resistant venetoclax-based therapy^62^. Given the results from our analysis of transcription factor activity in AML patients, we wanted to understand the extent to which *FOXO1* knockdown could sensitize monocytic AML to venetoclax treatment. We used CRISPR-Cas9 to knock-out *FOXO1* in THP-1 cells, a cell line model of monocytic leukemia (Supplementary Figure 5f-h). Both *FOXO1* CRISPR guides significantly increased venetoclax sensitivity in this cell line model of monocytic AML, suggesting *FOXO1* is an important mediator of venetoclax sensitivity (Figure 5d, e). These findings were consistent with the predictions from Priori and Norm WMEAN and demonstrate how Priori can be used to detect transcription factor mediators of drug resistance.

## DISCUSSION

We have developed Priori, a computational algorithm that infers transcription factor activity using prior biological knowledge. We demonstrated that Priori detects perturbed transcription factors with higher sensitivity and specificity than other methods. Our analyses show that this improved performance is due to assessment of the direction and impact of transcription factor regulation on their target genes. Using Priori, we identified FOXA1 activity as a potential regulator of survival in BIDC and nominated important downstream targets that may contribute to this survival difference. Moreover, we used Priori to nominate transcription factor regulators of drug sensitivity in AML. We found that predicted FOXO1 activity by Priori is highly associated with venetoclax resistance. We validated these findings in a cell line model of monocytic AML, which is resistant to venetoclax^62^.

Priori leverages the Pathway Commons resource to identify known gene regulatory networks in gene expression data. While using the relationships from Pathway Commons enables Priori to ground its findings in peer-reviewed literature, this may limit the discovery of novel regulatory relationships. Analyses from other groups suggest that prior-based method tend to replicate prior information^63^. Indeed, the analysis of BIDC patient samples revealed that Priori was able to identify known transcription factor drivers of BIDC. However, our findings also showed that Priori detected a relationship between FOXA1 activity and BIDC survival. While it has been shown that high *FOXA1* expression is associated with improved outcomes in patients with estrogen receptor-positive disease, these findings reveal a novel relationship between FOXA1 activity in estrogen receptor-positive and -negative disease^64–66^. Our analyses provide a novel mechanistic hypothesis that is ready for experimental investigation.

Pathway Commons integrates publicly available RNA, DNA, and protein data sourced from a variety of tissue types. However, Pathway Commons is not designed to curate tissue-specific transcription factor gene regulatory networks. Cell context influences regulatory interactions between transcription factors and their downstream target genes^2,67^. The single-gene perturbation gene expression dataset that we used to evaluate the transcription factor activity methods were performed in numerous cell types. The size of this dataset precluded a definitive evaluation of tissue-context as a determinant of method performance. However, we designed Priori to allow researchers to include their own regulatory networks for context-specific evaluation of their experiments. Overall, Priori should generate robust predictions that are generalizable across many cellular contexts.

Collectively, Priori, identifies aberrant transcription factor activity with high sensitivity and specificity. Priori is therefore a useful tool for the identification of important regulators of disease pathogenesis or drug sensitivity.

## ACKNOWLEDGEMENTS

We would like to thank all of the patients for their precious time and donation of samples supporting this research. We appreciate the OHSU core facilities ExaCloud Cluster Computational Resource and the Advanced Computing Center for their assistance. We would also like to thank Michal R. Grzadkowski for his advice in the early stages of Priori’s development.

## METHODS

### Priori Algorithm

#### Pre-Processing

Priori normalizes and scales the input RNA-seq data prior to downstream analysis. Priori first filters out counts with a standard deviation less than 0.1 (controlled by the *thresh_filter* parameter). Priori linearly shifts the remaining counts by the minimum value and then log_2_ normalizes them (*x*_*gene*_). Priori then scales the normalized counts (*z*_*gene*_) using one of four methods: “standard”

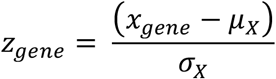

(where the mean and standard deviation of all normalized counts are indicated by *μ*_*x*_and *σ*_*x*_, respectively), “robust”

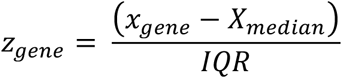

(where the median normalized counts is indicated by *X*_*median*_ and the interquartile range is indicated by *IQR*), “minmax”

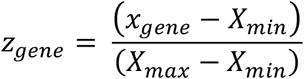

(where the minimum and maximum normalized counts are indicated by *X*_*min*_ and *X*_*max*_, respectively), or “quant” where the normalized counts are scaled using the inverse of the cumulative distribution function, *F(x)*^68^.

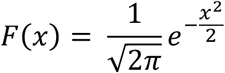

Priori defaults to the “standard” scaling function.

#### Network

Priori uses known gene regulatory networks to predict transcription factor activity from RNA-seq data. By default, Priori extracts regulatory relationships from the Pathway Commons database to generate activity scores^37,38^. Users can also generate Priori scores using other gene regulatory networks with the *regulon* parameter. The user-defined network must specify the transcription factor (*Regulator*) and their downstream target genes (*Target*). Pathway relationships in Pathway Commons are represented with the BioPAX language^37,69^. BioPAX abstracts major pathway relationships, including metabolic and signaling pathways, gene regulatory networks, and genetic and molecular interactions into a standardized format. However, BioPAX representations of gene regulatory networks cannot be interpreted directly. In order to identify the gene regulatory networks encoded in Pathway Commons, we extracted transcription factors and their primary targets using Patterns^70^. Using the extracted network, Priori removes transcription factors with less than 15 downstream targets. Users can control the number of targets with the *regulon_size* parameter.

#### Activity Scores

Once the network is prepared, Priori generates an activity score for each transcription factor in an RNA-seq dataset. To calculate the activity score for a transcription factor (*TF*_*i*_), Priori first calculates weights for its target genes 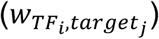. The target gene weight is the product of the F-statistic 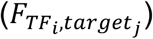 and the Spearman correlation coefficient 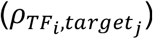:

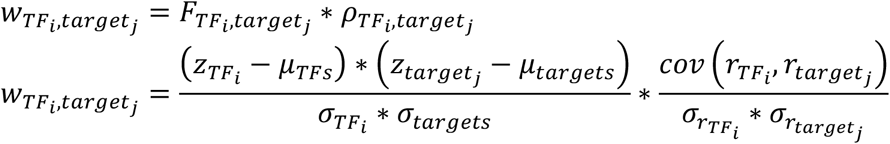

where *i* represents the range of transcription factors in a dataset, *j* represents the range of target genes for a given transcription factor *TF*_*i*_, and *r* represents the rank of the scaled counts relative to all features in the dataset. The non-negative, log_2_-normalized counts are used to calculate the F-statistic.

Priori first uses the transcription factor-target gene weights to determine the direction of regulation. Priori defines *k* down-regulated targets among all *j* target genes as those with 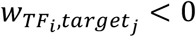. Priori also identifies *l* up-regulated targets among all *j* target genes as those with 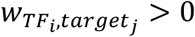. Priori then ranks the scaled target counts for each transcription factor grouped by their direction of regulation (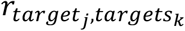 or 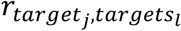). Priori weighs the ranks of the scaled counts using the target gene weight 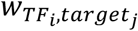:

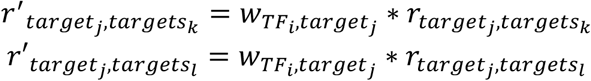

The resulting activity score for a given transcription factor 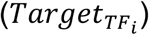 is the summation of the weighted ranks:

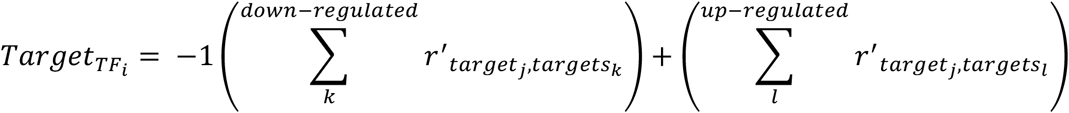

where downregulated genes are scaled by −1 prior to summation. Finally, the activity scores for each transcription factor are z-transformed relative to all other transcription factors and then again to all other samples in the dataset.

### Benchmarking Workflow

#### Pre-Processing

The decoupleR benchmarking workflow has been previously described^39^. The normalized RNA-seq data was linearly shifted by the minimum value so all values were non-negative^40^.

#### Transcription factor activity scores and p-values

With the decoupleR workflow (version 2.0.0), we generated transcription factor activity scores for 11 methods including Priori: AUCell (version 1.16.0), FGSEA (version 1.20.0), GSVA (version 1.42.0), MDT, ORA, UDT, ULM, VIPER (version 1.28.0), WMEAN, and WSUM. decoupleR also generated normalized transcription factor activity scores for FGSEA, WMEAN, and WSUM as well as corrected scores for WMEAN and WSUM^39^. We generated transcription factor activity scores for each method using the default parameters. We also calculated p-values for the Priori activity scores using a Student’s two-sided t-test with an FDR post-test correction.

#### Area under the receiver operating characteristic and precision recall curves

Using the decoupleRBench package (version 0.1.0), we generated AUROC and AUPRC values. Briefly, we ranked the absolute value of the activity scores from the decoupleR workflow for each experiment^39^. The activity scores were ranked separately for each method. We determined whether the perturbed transcription factors were among the top “n” scores. “n” was defined as the number of unique perturbed transcription factors in the dataset. As the number of unperturbed transcription factors in the dataset substantially outnumbered the perturbed factors, we deployed a down-sampling strategy to compare an equal number. We calculated AUROC and AUPRC values for 100 down-sampling permutations.

#### ARACNe Network

In order to better understand whether Priori’s performance advantages depend on the design of its input transcription factor network, we ran generated VIPER activity scores using transcriptional relationships from TCGA ARACNe networks and an alternative network using ARACNe-AP^25^. We downloaded the TCGA ARACNe networks using the arcane.networks package (version 1.18.0). For the ARACNe-AP network, we computed it from the Holland et al. perturbation dataset^40^. ARACNe-AP requires a list of transcription factors in order to generate a gene regulatory network. We used a list of transcription factors from the Alvarez et al. 2016 publication that was provided by Dr. Mariano Alvarez on September 25, 2019^26^. We excluded transcription factors that were not present in the Holland et al. dataset, resulting in an input list of 1,726 transcription factors. We ran ARACNe-AP (version 1.0, created with java 1.8.0_171-b11) with 100 bootstraps, --*p-value = 1E-8*, and --*random seeds = TRUE*^25^. The consolidated interactome included 1,726 transcription factors and 302,444 interactions.

### Data Analysis

#### Benchmarking Workflow

Custom scripts were used to evaluate the results of the decoupleR benchmarking workflow. In order to compare the scores across the different methods, we z-transformed the transcription factor activity scores. We calculated the Spearman correlation between the activity scores and p-values of each method.

#### TCGA BRCA

The normalized gene expression counts and metadata for the Breast Invasive Carcinoma (TCGA, PanCancer Atlas) study were downloaded from cBioPortal https://www.cbioportal.org^43,71^. Priori scores were generated from the normalized gene expression counts linearly shifted by the minimum value. VIPER scores were generated using the ARACNe BRCA network, *--pleiotropy = TRUE*, and *--eset*.*filter* = FALSE^25,26^. Norm WMEAN scores were generated using Pathway Commons transcriptional relationships, *--times = 100, --sparse = TRUE*, and *--randomize_type = rows*^39^. ORA scores were generated using Pathway Commons transcriptional relationships, *--n_up = 300, --n_down = 300, --n_background = 20000*, and *--with_ties = TRUE*. Custom scripts were used to exclude patients without basal or luminal BIDC. The Seurat package was used to perform dimensional reduction on the Priori scores by PCA and UMAP^72^. Survival data was analyzed for significance using a log-rank Mantel-Cox test. We assigned patients to “High” or “Low” groups depending on whether they were among the top 90% or bottom 10% of the value of interest. Differential gene expression networks were generated from the normalized gene expression data using CausalPath^52^. An FDR threshold of 0.001 was used to evaluate significant relationships following 100 permutations.

#### Beat AML

Priori scores were generated from the normalized gene expression counts of baseline AML patient samples from the Beat AML cohort^58,59^. These counts were also linearly shifted by the minimum value. VIPER scores were generated using the ARACNe-AP network LAML network, *--pleiotropy = TRUE*, and *--eset*.*filter* = FALSE^25,26^. For the ARACNe-AP network, we computed it from the normalized RNA-seq data from Beat AML^58,59^. ARACNe-AP requires a list of transcription factors in order to generate a gene regulatory network. We used a list of transcription factors from the Alvarez et al. 2016 publication that was provided by Dr. Mariano Alvarez on September 25, 2019^26^. We excluded transcription factors that were not present in the Holland et al. dataset, resulting in an input list of 1,408 transcription factors. We ran ARACNe-AP (version 1.0, created with java 1.8.0_171-b11) with 100 bootstraps, --*p-value = 1E-8*, and --*random seeds = TRUE*^25^. The consolidated interactome included 1,402 transcription factors and 320,632 interactions. Norm WMEAN scores were generated using Pathway Commons transcriptional relationships, *--times = 100, --sparse = TRUE*, and *--randomize_type = rows*^39^. ORA scores were generated using Pathway Commons transcriptional relationships, *--n_up = 300, --n_down = 300, --n_background = 20000*, and *--with_ties = TRUE*. Custom scripts were used to exclude patients without a diagnosis of AML or those with a prior myeloproliferative neoplasm. Priori scores and single-inhibitor drug AUC values on the same patient sample were evaluated using a Spearman correlation. Significant correlations were those with an FDR < 0.05.

### Cell Culture

#### Cell Lines

THP-1 cells (DSMZ) were cultured in RPMI (Gibco) supplemented with 10% fetal bovine serum (FBS, HyClone), 2 mM GlutaMAX (Gibco), 100 units/mL Penicillin, 100 μg/mL Streptomycin (Gibco), and 0.05 mM 2-Mercaptoethanol (Sigma Aldrich). All cells were cultured at 5% CO_2_ and 37°C. Cell lines were tested monthly for mycoplasma contamination.

#### CRISPR

Two *FOXO1* CRISPR guide RNAs in a pLentiCRISPR v2 backbone were obtained from GenScript^73^. The target sequence for guide RNA (gRNA) 1 is GCTCGTCCCGCCGCAACGCG and the sequence for gRNA 2 is ACAGGTTGCCCCACGCGTTG. Additionally, a non-targeting CRISPR guide RNA (target sequence CCTGGGTTAGAGCTACCGCA) generated by scrambling the target sequence of LentiCRISPRv2-ACTB-C1 in a pLentiCRISPR v2 backbone was obtained from Addgene (catalog #169795). Lentivirus was produced by transfecting Lenti-X 293T cells (Clontech) with the SMARTvector transfer plasmid and packaging/pseudotyping plasmids. psPAX2 was a gift from Didier Trono (Addgene plasmid #12260; http://n2t.net/addgene:12260; RRID:Addgene_12260). The supernatant containing lentivirus was collected after 48 hours of culture and filtered with a 0.45 μm filter. THP-1 cells were transduced with virus via spinnoculation in the presence of polybrene. Transduced cells were selected with 1 μg/mL puromycin to produce a stable cell line.

#### CRISPR Validation

*FOXO1* knockdown was validated using Tracking of Indels by Decomposition (TIDE)^74^. Briefly, cellular DNA was PCR-amplified using primers upstream (sequence AAGTAGGGCACGCTCTTGAC) and downstream (sequence CGTTCCCCCAAATCTCGGAC) of the *FOXO1* gRNA target sequences. The primers were designed in Geneious Prime and synthesized by Integrated DNA Technologies. Paramagnetic beads were used to purify the PCR DNA fragments (MagBio Ref #AC-60001) and subsequently sequenced by EuroFins. Inference of CRISPR Edits (ICE) was performed using the Synthego web tool (https://ice.synthego.com/#/).

#### Drug Sensitivity Assay

Cells were cultured for 72 hours along a 7-point dose curve with venetoclax. Cell viability was assessed by CellTiter Aqueous colorimetric assay.

### Availability of Data and Materials

Priori is free to use and is publicly accessible on GitHub (https://github.com/ohsu-comp-bio/regulon-enrichment). The forked versions of decoupleR (https://github.com/JEstabrook/decoupleR) and the benchmarking workflow (https://github.com/ohsu-comp-bio/decoupleRBench) were used in this study can be accessed on GitHub. The benchmark analysis pipeline is freely available on GitHub^75,76^ (https://github.com/ohsu-comp-bio/decoupler_workflow). The single-gene perturbation

RNA-seq data from *Holland et al*. analyzed in this study is accessible at: https://zenodo.org/record/3564179^40^. The TCGA BRCA RNA-seq data from BIDC patients analyzed during the current study is available on cBioPortal^43,71^. The Beat AML RNA-seq data from AML patients evaluated in this work is available on Vizome^58,59^.

## AUTHOR CONTRIBUTIONS

**J Estabrook**: conceptualization, software, formal analysis, validation, investigation, visualization, methodology, writing-review and editing. **WM Yashar**: conceptualization, software, formal analysis, validation, investigation, visualization, methodology, writing-original draft, writing-review and editing. **H Holly**: conceptualization, formal analysis, investigation, writing-review and editing. **J Somers**: formal analysis, investigation, writing-review and editing. **O Nikolova**: conceptualization, investigation, writing-review and editing. **Ö. Babur**: conceptualization, investigation, writing-review and editing. **TP Braun**: conceptualization, resources, supervision, funding acquisition, validation, investigation, writing-review and editing. **E Demir**: conceptualization, resources, formal analysis, supervision, funding acquisition, validation, investigation, visualization, methodology, writing-review and editing. The co-first authors may identify themselves as lead authors in their respective CVs.

## SUPPLEMENTAL FIGURES

**Supplementary Figure 1:**
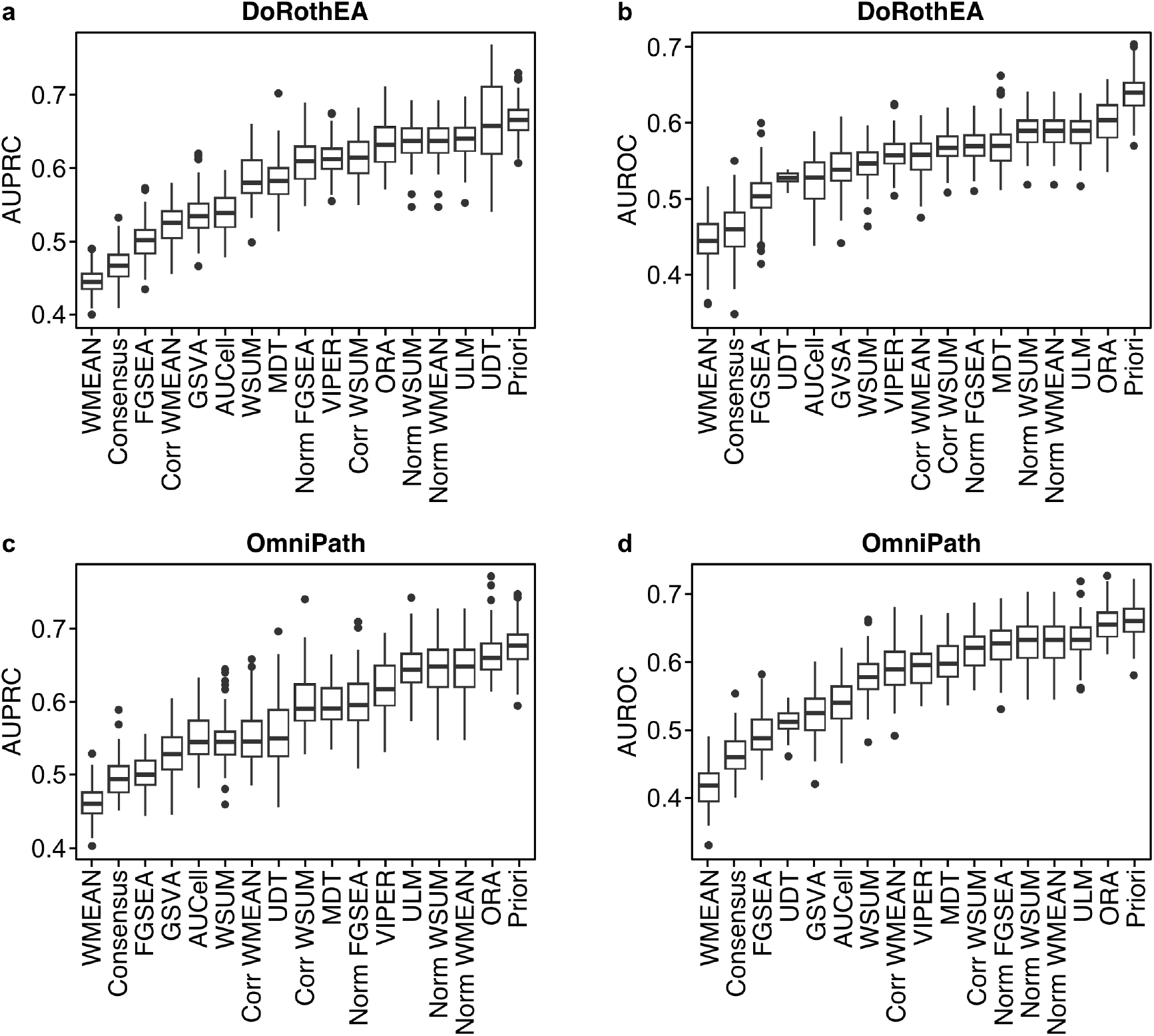
Priori demonstrated improved sensitivity and specificity when using transcriptional relationships from DoRoTHEA or OmniPath. a, b. Activity scores were generated using DoRothEA transcriptional relationships. The distribution of (a) AUPRC and (b) AUROC values across the 100 down-sampling permutations. c, d. Activity scores were generated using OmniPath transcriptional relationships. The distribution of (c) AUPRC and (d) AUROC values across the 100 down-sampling permutations.

**Supplementary Figure 2:**
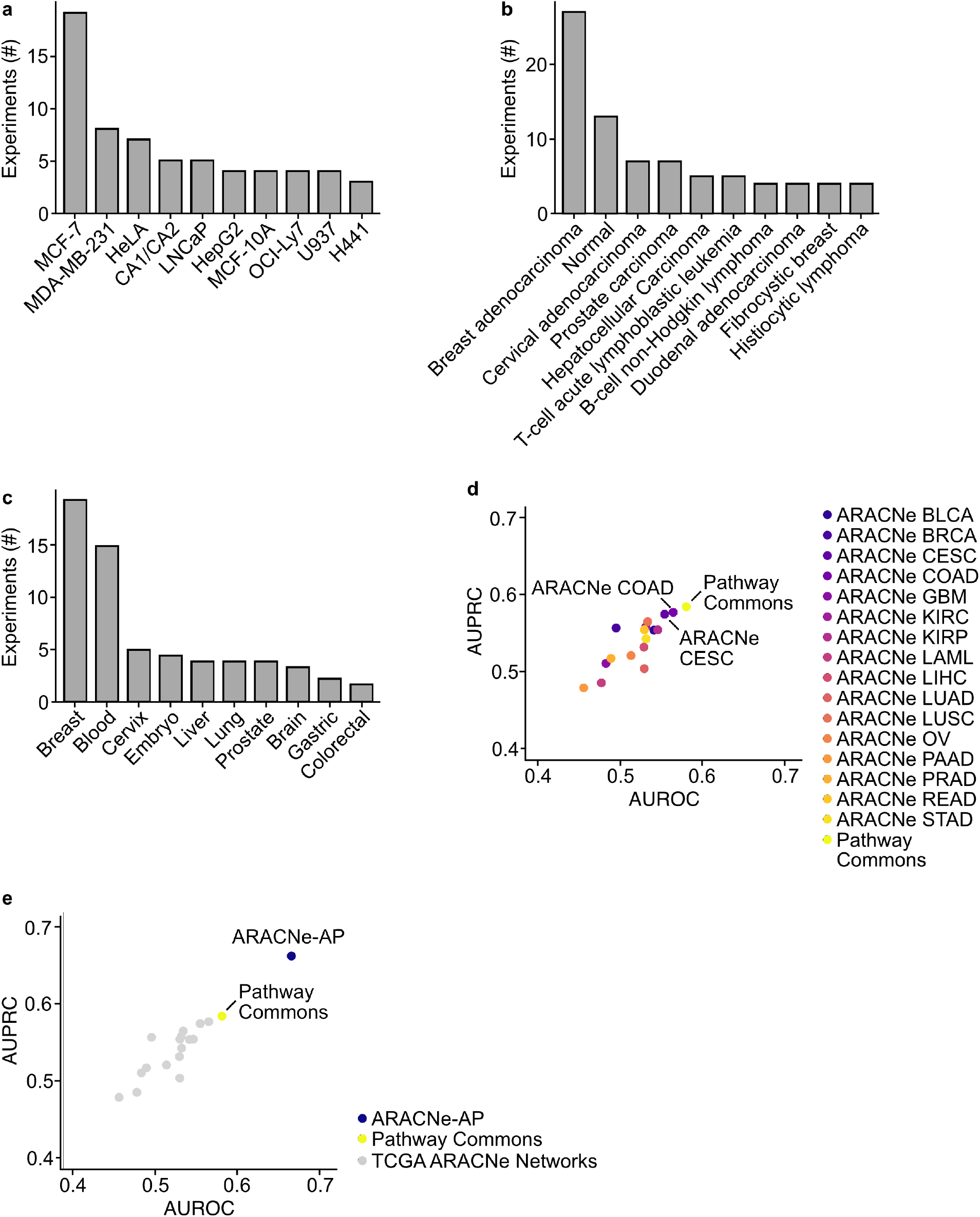
ARACNe-generated networks are important for VIPER performance. a-c. Top 10 represented (a) cell lines and their associated (b) diseases and (c) organ sites used in the perturbation dataset. d, e. VIPER activity scores were generated using transcriptional relationships from (d) TCGA ARACNe networks or (e) an ARACNe-AP network. The ARACNe-AP interactome was computed from the perturbation dataset. Mean AUPRC and AUROC values across the 100 down-sampling permutations for each network.

**Supplementary Figure 3:**
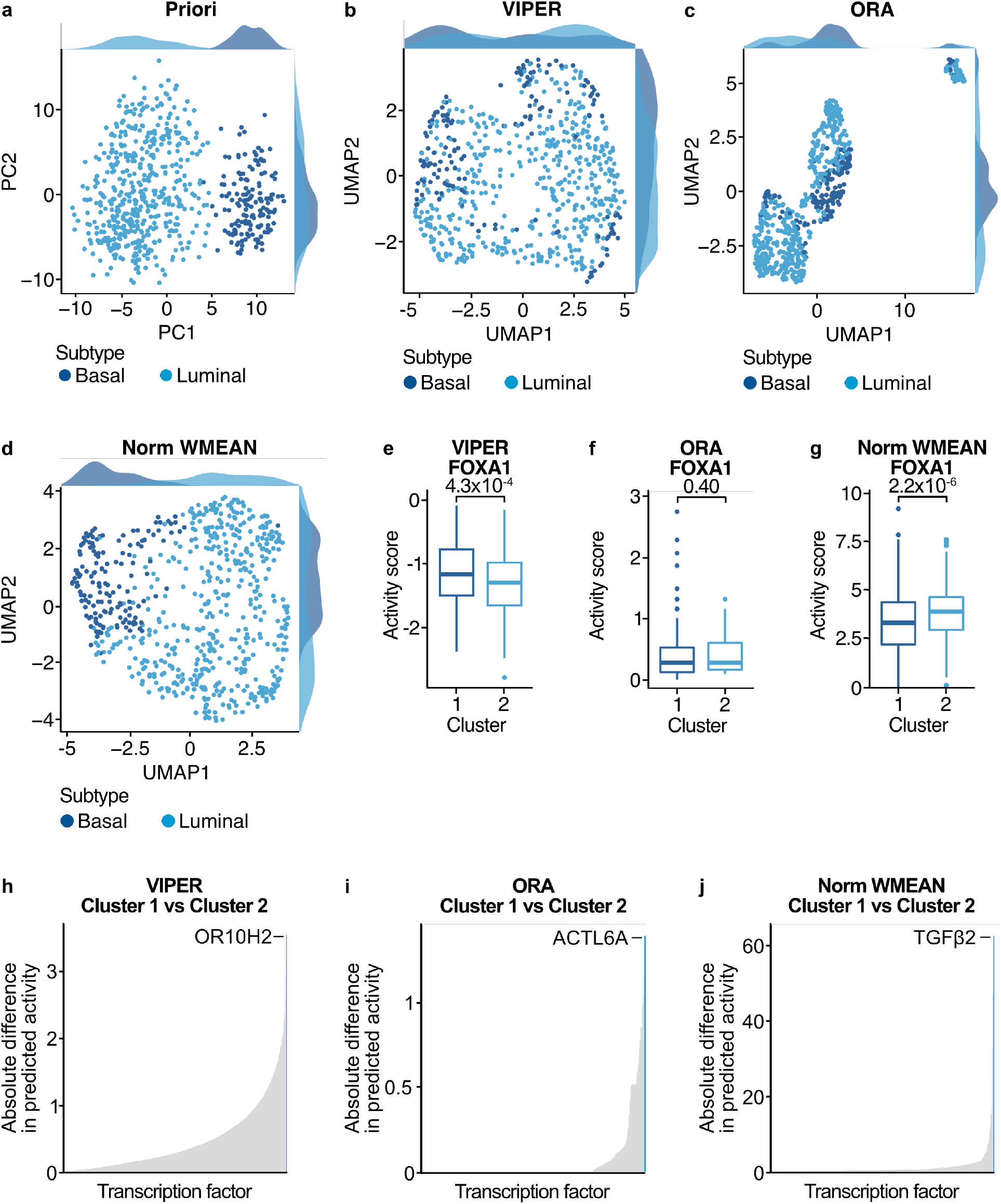
VIPER, ORA, and Norm WMEAN nominated distinct transcription factor regulators of BIDC pathogenesis. a. Priori scores were generated from RNA-seq of 637 patients with BIDC. PCA dimensional reduction and projection of Priori. Dots are colored by the breast cancer molecular subtype. b-d. Activity scores were generated using (b) VIPER, (c) ORA, and (d) Norm WMEAN from RNA-seq of 637 patients with BIDC. UMAP dimensional reduction and projection of activity scores. Dots are colored by the breast cancer molecular subtype. e-g. Unsupervised hierarchical clustering was performed using activity scores in (b-d). Distribution of FOXA1 (e) VIPER, (f) ORA, and (g) Norm WMEAN scores among patients in clusters 1 and 2 (defined separately for each method). h-j. Mean absolute difference of (h) VIPER, (i) ORA, and (j) Norm WMEAN activity scores from patients in clusters 1 and 2 (defined separately for each method). Transcription factor with the greatest absolute difference in activity scores between the two clusters is highlighted.

**Supplementary Figure 4:**
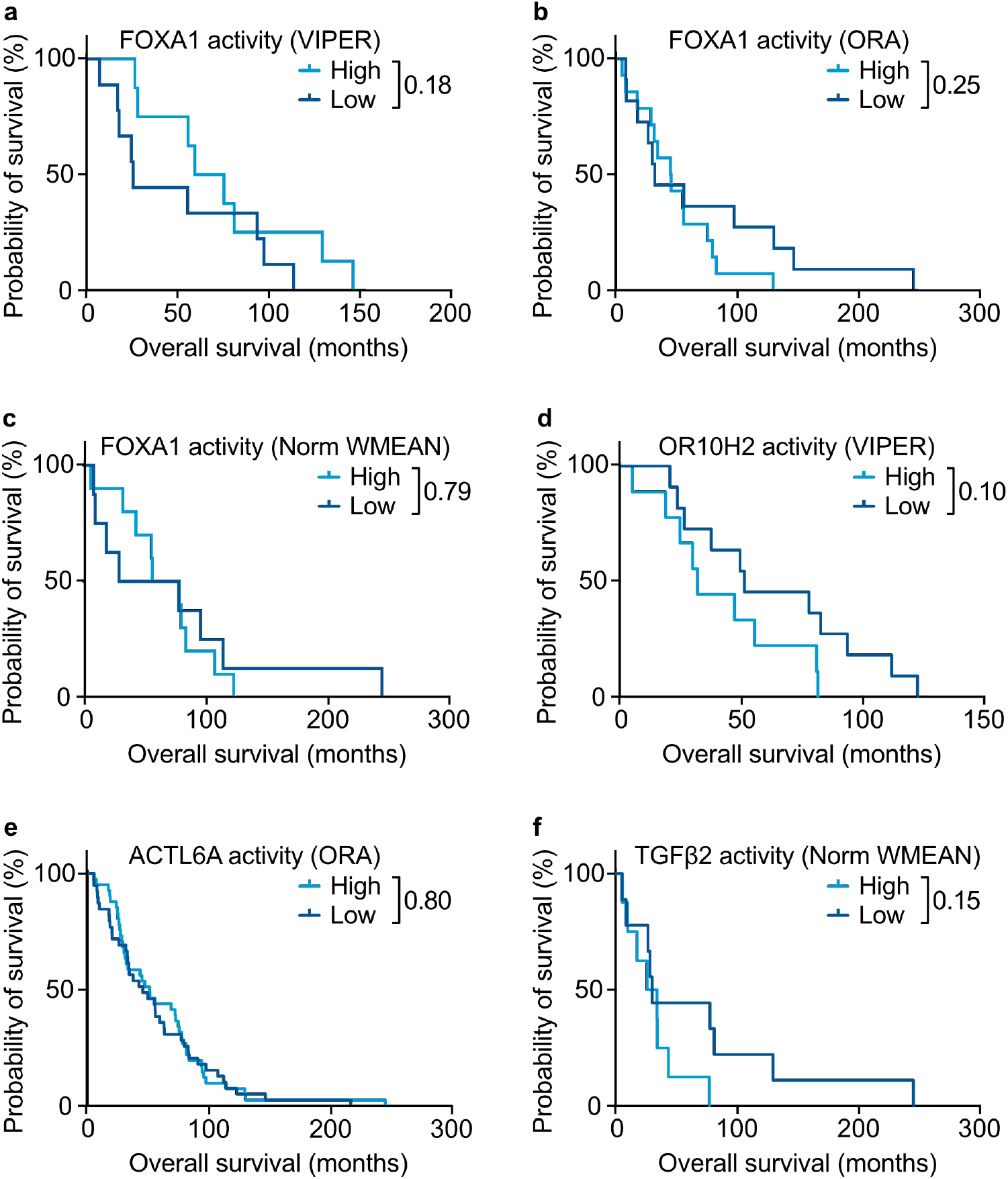
VIPER, ORA, and Norm WMEAN FOXA1 scores do not identify survival differences among patients with BIDC. a-f. Kaplan-Meier survival analysis of patients grouped by (a) FOXA1 VIPER scores, (b) FOXA1 ORA scores, (c) FOXA1 Norm WMEAN scores, (d) OR10H2 VIPER scores, (e) ACTL6A ORA scores, or (f) TGFβ2 Norm WMEAN scores. Patients among the top 90% of scores were grouped into “High” and those in the bottom 10% were grouped into “Low”. Statistical significance was determined by a log-rank Mantel-Cox test.

**Supplementary Figure 5:**
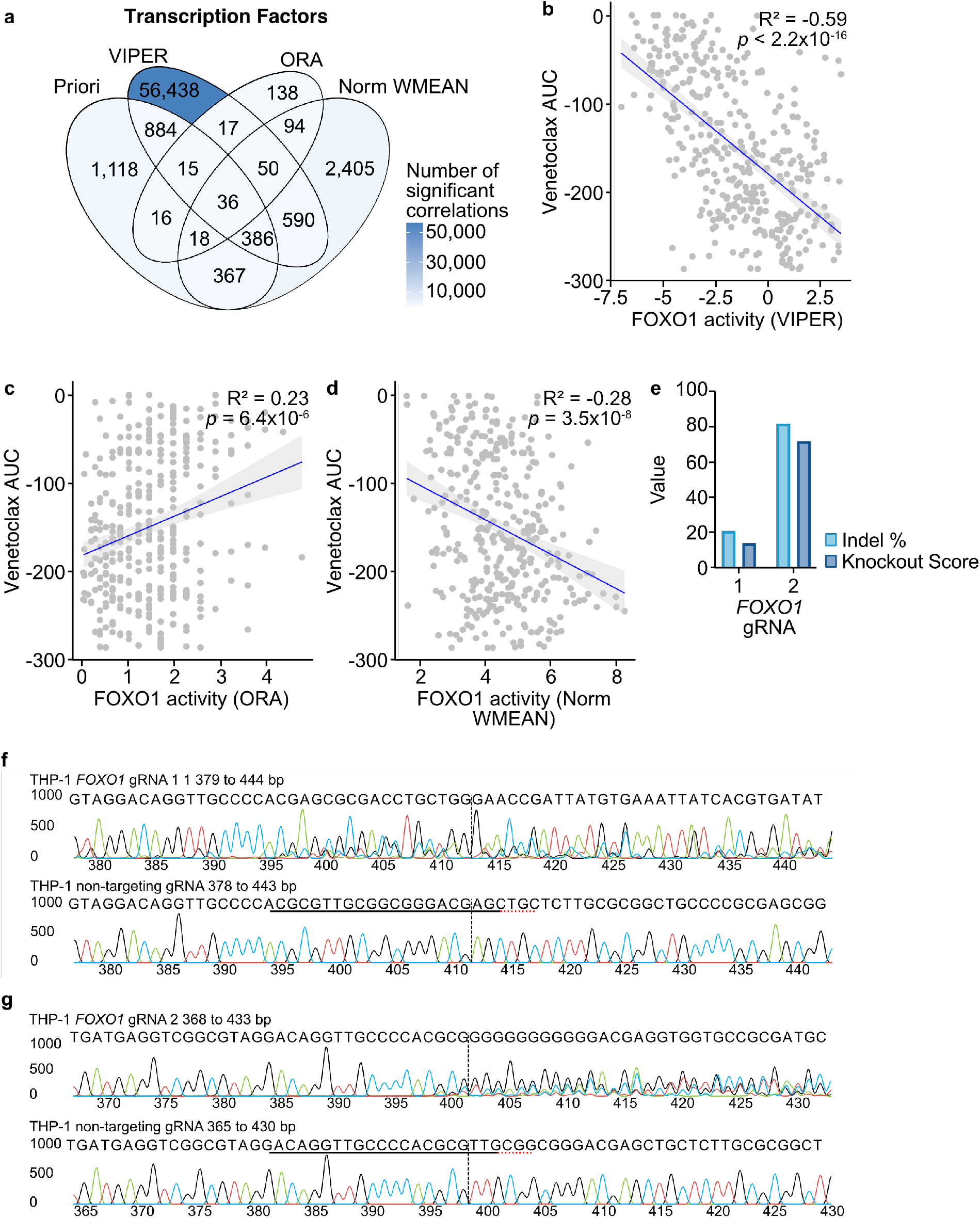
Validation of *FOXO1* knockdown by CRISPR. a. Activity scores were generated from RNA-seq of 859 patients with AML using VIPER, ORA, and Norm WMEAN. Activity scores for each method and *ex vivo* drug response AUC data were evaluated using Spearman correlation. Statistical significance was determined using the Spearman correlation p-value with an FDR post-test correction. Overlap of significant activity score-inhibitor correlations for each method (including the significant Priori correlations from Figure 5a). b-d. Spearman correlation of venetoclax AUC and FOXO1 (b) VIPER, (c) ORA, or (d) Norm WMEAN scores. Statistical significance was determined using the Spearman correlation p-value with an FDR post-test correction. e. THP-1 cells were transduced with lentiviral particles harboring expression cassettes for hSpCas9 as well as a non-targeting or *FOXO1* gRNA. Cells were evaluated by the percentage of indels and knockout score, which is the proportion of indels that are a frameshift or are greater than 21 bp in length. f, g. Sanger sequencing traces showing edited and control (cells with non-targeting gRNA) sequences in the region around the guide sequence. The horizontal black underlined region represents the guide sequence. The horizontal red underline represents the PAM site. The vertical black dotted line represents the cut site.

## REFERENCES

1. Bushweller, J. H. Targeting transcription factors in cancer — from undruggable to reality. Nat Rev Cancer 19, 611–624 (2019).

2. Lee, T. I. & Young, R. A. Transcriptional Regulation and Its Misregulation in Disease. Cell 152, 1237–1251 (2013).

3. Fuda, N. J., Ardehali, M. B. & Lis, J. T. Defining mechanisms that regulate RNA polymerase II transcription in vivo. Nature 461, 186–192 (2009).

4. Spitz, F. & Furlong, E. E. M. Transcription factors: from enhancer binding to developmental control. Nat Rev Genet 13, 613–626 (2012).

5. Sanda, T. et al. Core Transcriptional Regulatory Circuit Controlled by the TAL1 Complex in Human T Cell Acute Lymphoblastic Leukemia. Cancer Cell 22, 209–221 (2012).

6. Lin, C. Y. et al. Transcriptional Amplification in Tumor Cells with Elevated c-Myc. Cell 151, 56–67 (2012).

7. Nie, Z. et al. c-Myc is a universal amplifier of expressed genes in lymphocytes and embryonic stem cells. Cell 151, 68–79 (2012).

8. Khatri, P., Sirota, M. & Butte, A. J. Ten Years of Pathway Analysis: Current Approaches and Outstanding Challenges. PLOS Computational Biology 8, e1002375 (2012).

9. Nguyen, T.-M., Shafi, A., Nguyen, T. & Draghici, S. Identifying significantly impacted pathways: a comprehensive review and assessment. Genome Biol 20, 203 (2019).

10. Essaghir, A. et al. Transcription factor regulation can be accurately predicted from the presence of target gene signatures in microarray gene expression data. Nucleic Acids Research 38, e120 (2010).

11. de Sousa Abreu, R., Penalva, L. O., Marcotte, E. M. & Vogel, C. Global signatures of protein and mRNA expression levels. Mol Biosyst 5, 1512–1526 (2009).

12. Vogel, C. & Marcotte, E. M. Insights into the regulation of protein abundance from proteomic and transcriptomic analyses. Nat Rev Genet 13, 227–232 (2012).

13. Koussounadis, A., Langdon, S. P., Um, I. H., Harrison, D. J. & Smith, V. A. Relationship between differentially expressed mRNA and mRNA-protein correlations in a xenograft model system. Sci Rep 5, 10775 (2015).

14. Kiełbasa, S. M. & Vingron, M. Transcriptional Autoregulatory Loops Are Highly Conserved in Vertebrate Evolution. PLOS ONE 3, e3210 (2008).

15. Benito, J., Zheng, H., Ng, F. S. & Hardin, P. E. Transcriptional feedback loop regulation, function and ontogeny in Drosophila. Cold Spring Harb Symp Quant Biol 72, 437–444 (2007).

16. Bornstein, C. et al. A negative feedback loop of transcription factors specifies alternative dendritic cell chromatin states. Mol Cell 56, 749–762 (2014).

17. Califano, A. & Alvarez, M. J. The recurrent architecture of tumour initiation, progression and drug sensitivity. Nat Rev Cancer 17, 116–130 (2017).

18. Vaquerizas, J. M., Kummerfeld, S. K., Teichmann, S. A. & Luscombe, N. M. A census of human transcription factors: function, expression and evolution. Nat Rev Genet 10, 252–263 (2009).

19. Teschendorff, A. E. & Wang, N. Improved detection of tumor suppressor events in single-cell RNA-Seq data. npj Genom. Med. 5, 1–14 (2020).

20. Korotkevich, G. et al. Fast gene set enrichment analysis. 060012 Preprint at https://doi.org/10.1101/060012 (2021).

21. Hänzelmann, S., Castelo, R. & Guinney, J. GSVA: gene set variation analysis for microarray and RNA-Seq data. BMC Bioinformatics 14, 1–15 (2013).

22. Hung, J.-H., Yang, T.-H., Hu, Z., Weng, Z. & DeLisi, C. Gene set enrichment analysis: performance evaluation and usage guidelines. Briefings in Bioinformatics 13, 281–291 (2012).

23. Aibar, S. et al. SCENIC: Single-cell regulatory network inference and clustering. Nat Methods 14, 1083–1086 (2017).

24. Margolin, A. A. et al. ARACNE: An Algorithm for the Reconstruction of Gene Regulatory Networks in a Mammalian Cellular Context. BMC Bioinformatics 7, 1–15 (2006).

25. Lachmann, A., Giorgi, F. M., Lopez, G. & Califano, A. ARACNe-AP: gene network reverse engineering through adaptive partitioning inference of mutual information. Bioinformatics 32, 2233–2235 (2016).

26. Alvarez, M. J. et al. Functional characterization of somatic mutations in cancer using network-based inference of protein activity. Nat Genet 48, 838–847 (2016).

27. Olsen, C. et al. Inference and validation of predictive gene networks from biomedical literature and gene expression data. Genomics 103, 329–336 (2014).

28. Zhu, M., Liu, C.-C. & Cheng, C. REACTIN: regulatory activity inference of transcription factors underlying human diseases with application to breast cancer. BMC Genomics 14, 504 (2013).

29. Gao, F., Foat, B. C. & Bussemaker, H. J. Defining transcriptional networks through integrative modeling of mRNA expression and transcription factor binding data. BMC Bioinformatics 5, 31 (2004).

30. Tarca, A. L. et al. A novel signaling pathway impact analysis. Bioinformatics 25, 75–82 (2009).

31. Vaske, C. J. et al. Inference of patient-specific pathway activities from multi-dimensional cancer genomics data using PARADIGM. Bioinformatics 26, i237–i245 (2010).

32. Babur, O., Demir, E., Gönen, M., Sander, C. & Dogrusoz, U. Discovering modulators of gene expression. Nucleic Acids Res 38, 5648–5656 (2010).

33. Walhout, A. J. M. What does biologically meaningful mean? A perspective on gene regulatory network validation. Genome Biol 12, 109 (2011).

34. Barbuti, R., Gori, R., Milazzo, P. & Nasti, L. A survey of gene regulatory networks modelling methods: from differential equations, to Boolean and qualitative bioinspired models. J Membr Comput 2, 207–226 (2020).

35. Fernald, G. H., Capriotti, E., Daneshjou, R., Karczewski, K. J. & Altman, R. B. Bioinformatics challenges for personalized medicine. Bioinformatics 27, 1741–1748 (2011).

36. Yngvadottir, B., Macarthur, D. G., Jin, H. & Tyler-Smith, C. The promise and reality of personal genomics. Genome Biol 10, 237 (2009).

37. Cerami, E. G. et al. Pathway Commons, a web resource for biological pathway data. Nucleic Acids Res 39, D685–D690 (2011).

38. Rodchenkov, I. et al. Pathway Commons 2019 Update: integration, analysis and exploration of pathway data. Nucleic Acids Res 48, D489–D497 (2020).

39. Badia-i-Mompel, P. et al. decoupleR: ensemble of computational methods to infer biological activities from omics data. Bioinformatics Advances 2, vbac016 (2022).

40. Holland, C. H. et al. Robustness and applicability of transcription factor and pathway analysis tools on single-cell RNA-seq data. Genome Biol 21, 36 (2020).

41. Garcia-Alonso, L., Holland, C. H., Ibrahim, M. M., Turei, D. & Saez-Rodriguez, J. Benchmark and integration of resources for the estimation of human transcription factor activities. Genome Res. 29, 1363–1375 (2019).

42. Türei, D. et al. Integrated intra- and intercellular signaling knowledge for multicellular omics analysis. Molecular Systems Biology 17, e9923 (2021).

43. Berger, A. C. et al. A Comprehensive Pan-Cancer Molecular Study of Gynecologic and Breast Cancers. Cancer Cell 33, 690–705.e9 (2018).

44. Sørlie, T. et al. Gene expression patterns of breast carcinomas distinguish tumor subclasses with clinical implications. Proc Natl Acad Sci U S A 98, 10869–10874 (2001).

45. Perou, C. M. et al. Distinctive gene expression patterns in human mammary epithelial cells and breast cancers. Proc Natl Acad Sci U S A 96, 9212–9217 (1999).

46. Brenton, J. D., Carey, L. A., Ahmed, A. A. & Caldas, C. Molecular classification and molecular forecasting of breast cancer: ready for clinical application? J Clin Oncol 23, 7350–7360 (2005).

47. Tamimi, R. M. et al. Comparison of molecular phenotypes of ductal carcinoma in situand invasive breast cancer. Breast Cancer Res 10, 1–9 (2008).

48. Sorlie, T. et al. Repeated observation of breast tumor subtypes in independent gene expression data sets. Proc Natl Acad Sci U S A 100, 8418–8423 (2003).

49. Kouros-Mehr, H. & Werb, Z. Candidate regulators of mammary branching morphogenesis identified by genome-wide transcript analysis. Dev Dyn 235, 3404–3412 (2006).

50. Kouros-Mehr, H., Slorach, E. M., Sternlicht, M. D. & Werb, Z. GATA-3 Maintains the Differentiation of the Luminal Cell Fate in the Mammary Gland. Cell 127, 1041–1055 (2006).

51. Sotiriou, C. et al. Breast cancer classification and prognosis based on gene expression profiles from a population-based study. Proc Natl Acad Sci U S A 100, 10393–10398 (2003).

52. Babur, Ö. et al. Causal interactions from proteomic profiles: Molecular data meet pathway knowledge. PATTER 0, (2021).

53. Chen, X. et al. XBP1 promotes triple-negative breast cancer by controlling the HIF1α pathway. Nature 508, 103–107 (2014).

54. Garcia-Alonso, L. et al. Transcription Factor Activities Enhance Markers of Drug Sensitivity in Cancer. Cancer Research 78, 769–780 (2018).

55. Alessandrini, F., Pezzè, L., Menendez, D., Resnick, M. A. & Ciribilli, Y. ETV7-Mediated DNAJC15 Repression Leads to Doxorubicin Resistance in Breast Cancer Cells. Neoplasia 20, 857–870 (2018).

56. Neel, D. S. & Bivona, T. G. Resistance is futile: overcoming resistance to targeted therapies in lung adenocarcinoma. NPJ Precis Oncol 1, 3 (2017).

57. Matkar, S. et al. An Epigenetic Pathway Regulates Sensitivity of Breast Cancer Cells to HER2 Inhibition via FOXO/c-Myc Axis. Cancer Cell 28, 472–485 (2015).

58. Tyner, J. W. et al. Functional genomic landscape of acute myeloid leukaemia. Nature 562, 526–531 (2018).

59. Bottomly, D. et al. Integrative analysis of drug response and clinical outcome in acute myeloid leukemia. Cancer Cell 40, 850–864.e9 (2022).

60. Mihalyova, J. et al. Venetoclax: A new wave in hematooncology. Experimental Hematology 61, 10–25 (2018).

61. Brown, F. et al. PRMT5 Inhibition Promotes FOXO1 Tumor Suppressor Activity to Drive a Pro-Apoptotic Program That Creates Vulnerability to Combination Treatment with Venetoclax in Mantle Cell Lymphoma. Blood 138, 681 (2021).

62. Pei, S. et al. Monocytic Subclones Confer Resistance to Venetoclax-Based Therapy in Patients with Acute Myeloid Leukemia. Cancer Discov 10, 536–551 (2020).

63. Hawe, J. S. et al. Network reconstruction for trans acting genetic loci using multi-omics data and prior information. Genome Med 14, 1–21 (2022).

64. Perou, C. M. et al. Molecular portraits of human breast tumours. Nature 406, 747–752 (2000).

65. Lupien, M. et al. FoxA1 Translates Epigenetic Signatures into Enhancer-Driven Lineage-Specific Transcription. Cell 132, 958–970 (2008).

66. Hurtado, A., Holmes, K. A., Ross-Innes, C. S., Schmidt, D. & Carroll, J. S. FOXA1 is a key determinant of estrogen receptor function and endocrine response. Nat Genet 43, 27–33 (2011).

67. MacNeil, L. T. et al. Transcription Factor Activity Mapping of a Tissue-Specific In Vivo Gene Regulatory Network. cels 1, 152–162 (2015).

68. Pedregosa, F. et al. Scikit-learn: Machine Learning in Python. Journal of Machine Learning Research 12, 2825–2830 (2011).

69. Demir, E. et al. BioPAX – A community standard for pathway data sharing. Nat Biotechnol 28, 935–942 (2010).

70. Babur, Ö. et al. Pattern search in BioPAX models. Bioinformatics 30, 139–140 (2014).

71. Cerami, E. et al. The cBio cancer genomics portal: an open platform for exploring multidimensional cancer genomics data. Cancer Discov 2, 401–404 (2012).

72. Hafemeister, C. & Satija, R. Normalization and variance stabilization of single-cell RNA-seq data using regularized negative binomial regression. Genome Biology 20, 296 (2019).

73. Sanjana, N. E., Shalem, O. & Zhang, F. Improved vectors and genome-wide libraries for CRISPR screening. Nat Methods 11, 783–784 (2014).

74. Brinkman, E. K., Chen, T., Amendola, M. & van Steensel, B. Easy quantitative assessment of genome editing by sequence trace decomposition. Nucleic Acids Research 42, e168 (2014).

75. Mölder, F. et al. Sustainable data analysis with Snakemake. Preprint at https://doi.org/10.12688/f1000research.29032.2 (2021).

76. Köster, J. & Rahmann, S. Snakemake—a scalable bioinformatics workflow engine. Bioinformatics 34, 3600–3600 (2018).

